# Interfacial Water Molecules Make RBD of SPIKE Protein and Human ACE2 to Stick Together

**DOI:** 10.1101/2020.06.15.152892

**Authors:** Ashish Malik, Dwarakanath Prahlad, Naveen Kulkarni, Abhijit Kayal

## Abstract

A novel coronavirus (SARS-CoV-2; COVID-19) that initially originates from Wuhan province in China has emerged as a global pandemic, an outbreak that started at the end of 2019 which claims 431,192 (Date: 15^th^ June 2020 (https://covid19.who.in) life till now. Since then scientists all over the world are engaged in developing new vaccines, antibodies, or drug molecules to combat this new threat. Here in this work, we performed an *in-silico* analysis on the protein-protein interactions between the receptor-binding (RBD) domain of viral SPIKE protein and human angiotensin-converting enzyme 2 (hACE2) receptor to highlight the key alteration that happened from SARS-CoV to SARS-CoV-2. We analyzed and compared the molecular differences between these two viruses by using various computational approaches such as binding affinity calculations, computational alanine, and molecular dynamics simulations. The binding affinity calculations show SARS-CoV-2 binds little more firmly to the hACE2 receptor than that of SARS-CoV. Analysis of simulation trajectories reveals that enhanced hydrophobic contacts or the van der Waals interaction play a major role in stabilizing the protein-protein interface. The major finding obtained from molecular dynamics simulations is that the RBD-ACE2 interface is populated with water molecules and interacts strongly with both RBD and ACE2 interfacial residues during the simulation periods. We also emphasize that the interfacial water molecules play a critical role in binding and maintaining the stability of the RBD/hACE2 complex. The water-mediated hydrogen bond by the bridge water molecules is crucial for stabilizing the RBD and ACE2 domains. The structural and dynamical features presented here may serve as a guide for developing new drug molecules, vaccines, or antibodies to combat the COVID-19 pandemic.

## 1. Introduction

The novel coronavirus (severe acute respiratory syndrome coronavirus 2, SARS-CoV-2) has emerged as a global pandemic by taking more than 431,192 (Date: 15^th^ June 2020 (https://covid19.who.in) life all over the world. The rapid spread and high mortality rate (3 to 5%)^1–3^ of this virus pose a serious global health emergency^4,5^. The general symptoms are headache, high fever, severe respiratory illness, pneumonia. So far no suitable vaccine or drug molecule is available to stop this virus from spreading. The new SARS-CoV-2 is characterized as a new member of the bat coronavirus genome and closely related to the severe acute respiratory syndrome coronavirus (SARS-CoV)^4^. SARS-CoV is a similar kind of virus that already created a pandemic in the year 2002. But the magnitude of fatalitity caused by SARS-CoV-2 surpasses all the previous SARS-like viruses in terms of cost to human life. Both the coronaviruses, SARS-CoV and SARS-CoV-2, use the homotrimer spike glycoprotein (comprising of S1 and S2 subunit) or SPIKE protein to bind the cellular receptors and induce the dissociation of S1 and S2 subunit. This also triggers a cascade of events such as the transition of S2 from a metastable prefusion state to a more stable post-fusion state, the fusion between a cell and viral membrane, etc^6–8^. S1 subunit contains the receptor-binding domain (RBD) that binds to the peptidase domain (PD) of human angiotensin-converting enzyme 2 (ACE2)^9^. The recognition of SPIKE protein with the ACE2 enzyme is the first step of the virus lifecycle and entry to the host cell. These are the reasons the CoV SPIKE protein is a key target for developing new vaccines, therapeutic antibodies, and new drug molecules. Recent in vitro studies also indicate that the RBD domain of the S1 subunit plays a key functional role for the binding of SARS-CoV by ACE2^10^. The crystal as well as cryo-EM structures of the complexes (SPIKE/hACE2) reveal key information about the interactions between the RBDs and the hACE2 receptor^6,9,11^. Superposition of RBD domains of SARS-CoV and SARS-CoV-2 shows very similar structural fold with a root mean square deviation (RMSD) of 0.68 Å of 139 pairs of Cα atoms^12^. Knowing that binding to the ACE2 receptor triggers the virial lifecycle, it is also important to understand how the binding affinity of SARS-CoV-2 differs from the SARS-CoV. Experimental results reveal that the binding affinity values for SARS-CoV and SARS-CoV-2 with hACE2 fall in a similar range^12^. Specifically, the equilibrium dissociation constant (K_*D*_) shows a slightly higher value (15.2 nm) of SARS-CoV-2 than that of SARS-CoV (15 nM)^13^. However, a recent publication shows a 20-fold increased binding affinity between trimeric SPIKE protein with ACE2 receptor of SARS-CoV-2 than that of SARS-CoV^6^. Nevertheless, it is now well established that SARS-CoV-2 has a slightly better binding affinity towards hACE2 than the SARS-CoV. The high binding affinity suggests that this novel coronavirus is evolved towards being a better binder to the same human ACE2 receptor. Thus, a comparison of both the RBD domains of SARS-CoV and SARS-CoV-2 is of utmost importance and it migh t give us a clue why SARS-CoV-2 is so deadlier than SARS-CoV.

Similar to the experiments, several computational studies are performed on the binding affinity of the RBDs of SARS-CoV and SARS-CoV-2 with hACE2^14,15^. Most of the earlier results are based on the homology modeled structure of the SARS-CoV-2, where the global fold of the SPIKE protein was correct but the atomic level scoring function for predicting the binding affinity was not accurate. Thus the earlier results predicted lower binding affinity of SARS-CoV-2^14,15^. This also implies that the side-chain packing of the modeled structure was not accurate enough. Recent binding free energy calculations performed on the crystal structure of RBD/hACE2 of SARS-CoV-2 predicts high binding affinity than SARS-CoV^16^. Another study uses Rosetta Interface Analyzer^17^ and concludes that both these viruses have a similar binding affinity^18^. Thus, from *in-silico* perspective, it is still not fully understood why SARS-CoV-2 is better binder than SARS-CoV-2 and the results are inconclusive. Hence, atomistic-level comparison is needed between these two viruses. A recent study found that the hydrogen bonding network and the hydrophobic interactions are major dominating forces that are responsible for enhanced binding in SARS-CoV-2^19^. Till now most of the studies focus on the search for new vaccines and drug molecules against COVID-19. There is very little attention paid to the atomistic label description and dynamics of the interfacial domain of SPIKE protein with the hACE2 receptor. The dynamic interactions between the RBD and the hACE2 may reveal some crucial points that are not accessible from only the crystal structures.

In this work, we first assess the atomistic level description of RBDs of SPIKE protein with hACE2 receptor from the crystal structures. We identify the key contact residues that are responsible for the binding of the RBD domain and hACE2. We also critically analyze the differences in the interfacial residues between the SARS-CoV and SARS-CoV-2. Further the binding affinity of these two domains of SARS-CoV and SARS-CoV-2. Next, all-atom molecular dynamics simulations of RBD domains of SPIKE protein complexed with the human ACE2 receptor of SARS-CoV and SARS-CoV-2 are performed to extract the dynamics of these complexes. We focus our study at the interfacial region of RBDs and hACE2 and provide an atomistic picture of the interactions. The importance of contact residues is also highlighted and a thorough comparison was made between the SARS-CoV and SARS-CoV-2. Till now, most of the *in-silico* work on SPIKE-ACE2 complex is done to design new antibodies or drug molecules, whereas little attention is paid to the atomistic level description and the dynamics of the SPIKE-ACE2 complex. Hence, our study focuses on the interfacial region of RBD and ACE2 domains. Our simulation results reveal that the interfacial water molecules play an important role regarding the binding of RBDs and ACE2 domain, particularly, the bridge water connecting the two domains of SPIKE protein and the hACE2.

## 2. Computational Details

The crystal structure of SARS-CoV-2-RBD/ACE2 (PDB ID: 6VW1) and SARS-CoV-RBD/ACE (PDB ID: 2AJF) are taken from the protein database and subjected to MD simulations. The two protein-protein complexes contain some notable structural elements for which special care was taken while preparing the system, like N-glycosylation of various asparagine residues, disulfide bridges between various pairs of cysteine, presence of zinc finger and other amino acids which require proper protonation states. The systems are prepared using the CHARMM-GUI web server^20^ and parameter files and equilibration files are taken from the server. All the simulations were performed using the software GROMACS version 2019.4^21,22^ using the CHARMM-36 forcefield^23^. Each system is solvated in TIP3P^24^ water and KCl salt were added to neutralize the systems to a salt concentration of 150 mM. Each system is solvated in a rectangular box of dimensions L_*x*_=138 Å, L_*y*_=138 Å, and L_*z*_=138 Å, the distance is appropriate to avoid all finite-size effects due to long-range electrostatic interactions among neighboring simulation boxes. Simulation for both the complexes is carried out at 310K and 1 atm pressure. To treat long-range electrostatic interactions, Particle Mesh Ewald (PME)^25^ method is used with a cut-off of 12 Å, and to constrain hydrogen bond, the LINCS algorithm^26^ was applied. Each system is minimized with 5000 steps of the steepest descent algorithm to eliminate the bad contacts in the system. After minimization equilibration simulation is performed for in NVT ensemble at 310 K. After NVT simulation, NPT equilibration is performed for 5 ns at 1 atm pressure, and the temperature is maintained at 310 K. During the equilibration period, the position restrained is applied to the heavy atoms of the protein. Velocity rescale thermostat and Parinnello-Rahman Barostat^27^ are applied to maintain the temperature and the pressure respectively. For the protein-protein binding affinity calculations, FoldX software^28^ is used. Different visualizing and trajectory analysis software packages UCSF Chimera^29^ and VMD (Visual Molecular Dynamics) ^30^ were used and QZyme Workbench™^31^ an in-house enzyme engineering platform was used for all calculations performed and discussed in results.

## 3. Results and Discussion

### 3.1. Structure and Energetics Comparison between SARS-CoV and SARS-CoV-2

Earlier published results highlighted the differences between the RBD domains of two SPIKE proteins. The SPIKE protein of SARS-CoV and SARS-CoV-2 share 75% sequence similarity and their structural fold are almost identical. Whereas the RBD domains have a sequence similarity of 73.7%^4^. **Figure 1** shows the superposition of SARS-CoV and SARS-CoV-2 and in the inset, we highlight the key residues that have changed from SARS-CoV to SARS-CoV-2. In the crystal structure of SARS-CoV and SARS-CoV-2, there are significant protein-protein contacts that can be seen and we identified some of the key residues that are important for the stabilization of SPIKE protein and hACE2 receptor. In **Figure S1** of **Supporting Information,** we displayed the residues that are in the interfacial region of the RBD domain bound to the ACE2 domain. With a distance cut-off 4 Å there are 16 residues of RBD that are in contact with 20 residues of ACE2 domain. A similar analysis of SARS-CoV-2 shows 18 residues of RBD that are in contact with 21 residues of the ACE2 receptor. Among the 21 ACE2 residues that are interacting with two RBDs, 17 are found to be common residues. Most of the interacting residues of the ACE2 receptor are located near the N-terminal helix. The above analysis manifests that the surface area of the interfacial domain of ACE2 and RBDS is almost the same. This also corroborates earlier findings where the surface area is 1,687 Å^2^ and 1,699 Å^2^ for SARS-CoV-2 and SARS-CoV^32^. Within this 4 Å cut-off there are 4 hydrophobic residues (Leu455, Phe456, Ala475, and Phe486) and 14 hydrophilic *(i.e.* not hydrophobic) residues (Arg408, Thr415, Tyr449, Tyr453, Gly476, Asn487, Tyr489, Gln493, Gly496, Gln498, Thr500, Asn501, Gly502, and Tyr505) are there near the interface of SPIKE protein of SARS-CoV-2. Whereas there are 3 hydrophobic residues (Leu443, Leu472, and Ile489) and 13 hydrophilic residues (Arg426, Tyr436, Tyr440, Tyr442, Asn473, Tyr475, Asn479, Gly482, Tyr484, Thr486, Thr487, Gly488, Tyr491) are present near the interface of SPIKE protein of SARS-CoV. It is interesting to note that the interface of RBD domain is enriched with tyrosine and glycine residues for SARS-CoV-2 (4 tyrosine and 3 glycine out of 14 residues) as well as SARS-CoV (6 tyrosine and 2 glycine out of 16 residues). Further, we counted the residues with a cut-off value of 8 Å to get an estimation of the buried residues of both the RBD domains. We found that 53 and 51 residues are coming in contact with the ACE2 domain for SARS-CoV and SARS-CoV-2 respectively (**Table S1, Supporting Information**). To see the effect of the interfacial residues on the overall binding affinity with hACE2, we performed computational alanine scanning on these interfacial residues to see the extent of the impact on the binding affinity due to alanine substitution.

**Figure 1.**
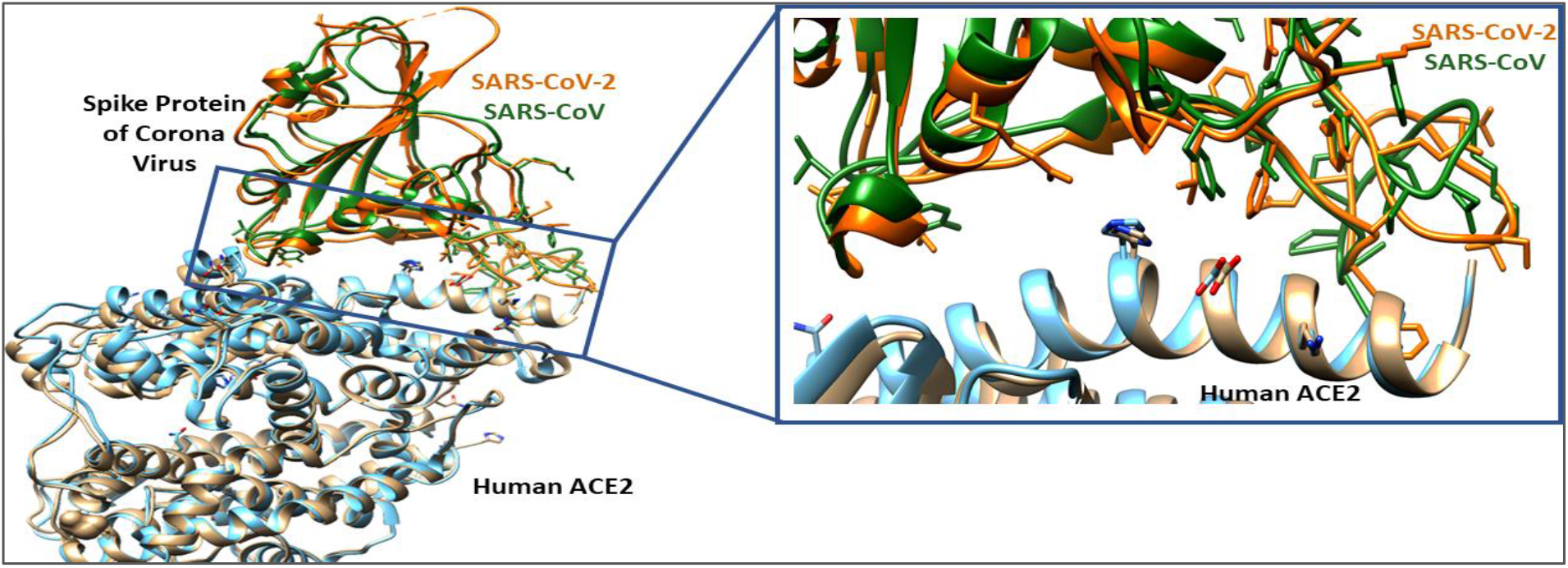
Structural alignment of RBD domain of SPIKE protein and human ACE2 receptor of SARS-CoV and SARS-CoV-2. For superposition, PDB ID (2AJF) and PDB ID (6VW1) are used for SARS-CoV and SARS-CoV-2. Inset we are showing the residues that are mutated from 2002 to 2019 in Licorice form.

Further, the crystal structure of SARS-CoV-2 has 13 hydrogen bonds between the RBD domain and the ACE2 domain. However, SARS-CoV has a lesser number of hydrogen bonds between the RBD and the ACE2 domain for SARS-CoV. It has only 5 hydrogen bonds between the two domains. As there are fewer hydrogen bonds in SARS-CoV, we expect that the RBD/ACE2 complex of SARS-CoV-2 is more stable than SARS-CoV. Further, we calculate the binding free energy of the SPIKE/hACE2 complex from the PRODIGY web server^33^ as well as from FoldX software^28^. The binding free energy is calculated on the crystal structures of the dimer and the values are −10.8 and −11.7 kcal/mol for SARS-CoV and SARS-CoV-2 respectively. While the FoldX calculations show binding energy −4.49 and −9.87 kcal/mol for SARS-CoV and SARS-CoV-2 respectively. Though the values obtained from PRODIGY and FoldX are not in the same range, the trend suggests that SARS-CoV-2 binds more effectively to hACE2 and evolved as a better binder to the human receptor.

### 3.2 In Silico-Alanine Scanning of Interfacial Residues of RBD Domain

As we discussed in the previous section about the interfacial residues of the RBD domain is critical for stabilizing the protein-protein complex, we performed computational alanine scanning on the contact residues of the RBD domain to highlight the importance of those residues. We mutated the interfacial residues to alanine and then calculate the binding free energy with the ACE2 domain. Alanine scanning is performed to identify the residues that make a significant contribution to the binding affinity with hACE2. The result reveals that (Figure 3) most of the residues lower the binding affinity when mutated with alanine except N501 where a slight increase in binding affinity is observed for SARS-CoV-2. A decrease in binding affinity suggests that both SARS-CoV-2 and SARS-CoV have optimized its RBD domain to bind with the human ACE2 receptor. From the alanine scanning results, we observe that Y449, L455, F486, Y498, G502, and V503 are found to be critical for stabilizing the overall complex. Compared to SARS-CoV-2, the alanine mutation doesn’t lower the binding affinity for all the residues of SARS-CoV. This result points to the fact that SARS-CoV-2 optimized its interfacial residues in such a manner that binding to hACE2 is more enhanced. In this context, we would like to discuss a few changes that happened between the SARS-CoV to SARS-CoV-2. Few mutations like Tyr484→Gln498, Thr487→Asn501, Tyr442→Leu455, Leu443→Phe456, Leu472→Phe486 Asn479→Gln493 show significant changes in binding affinity when the particular position is mutated with alanine. It is to be noted that Gln474 and Gly485 inserted near the interfacial domain of RBD of SARS-CoV-2. Though these two residues are not directly interacting with the ACE2, they form intramolecular hydrogen bonds two give the stability of the loop. Like Gln474 interacts with Gly476 and Gly484 interacts with Cys488. The alanine mutation of these two residues does not change the binding affinity significantly.

**Figure 2.**
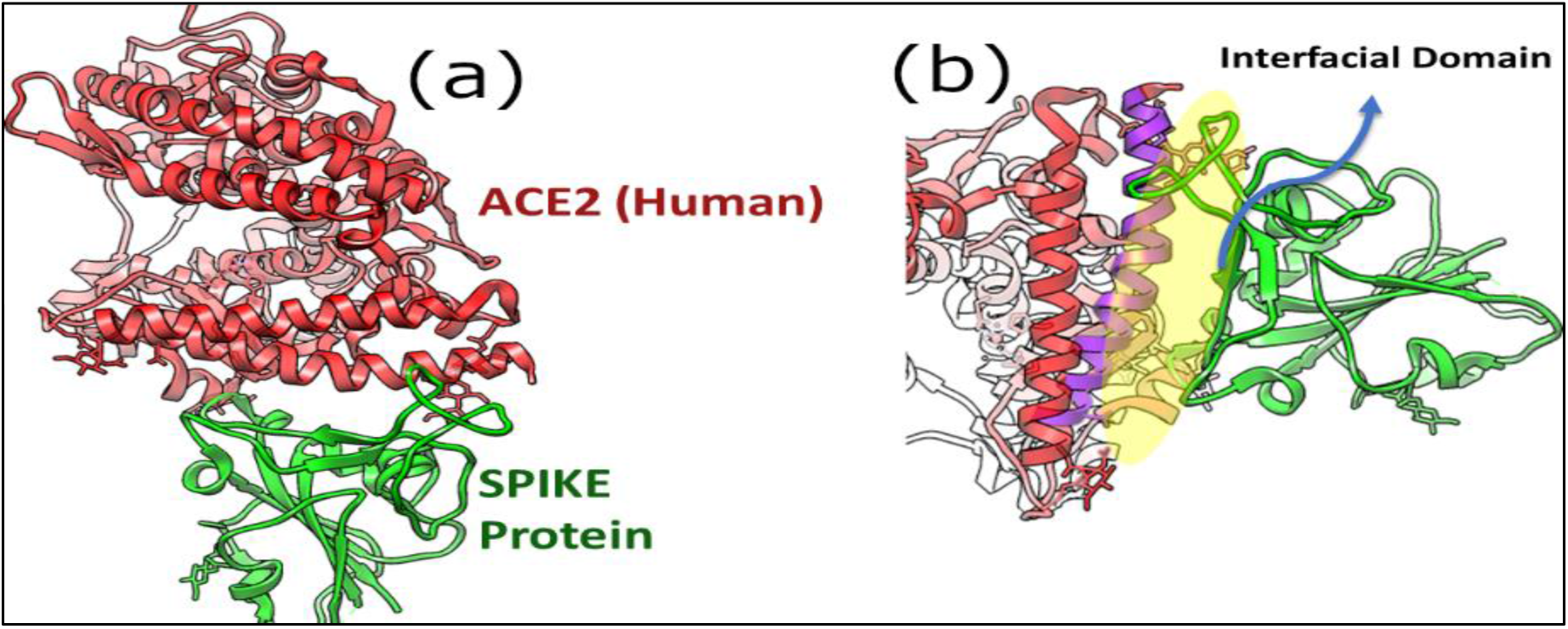
(a) Crystal structure of SARS-CoV-2 (PDB ID: 6VW1). (b) The yellow shaded area represents the space or the interfacial region in bewteen the ACE2 and the RBD domain of the SPIKE protein.

**Figure 3.**
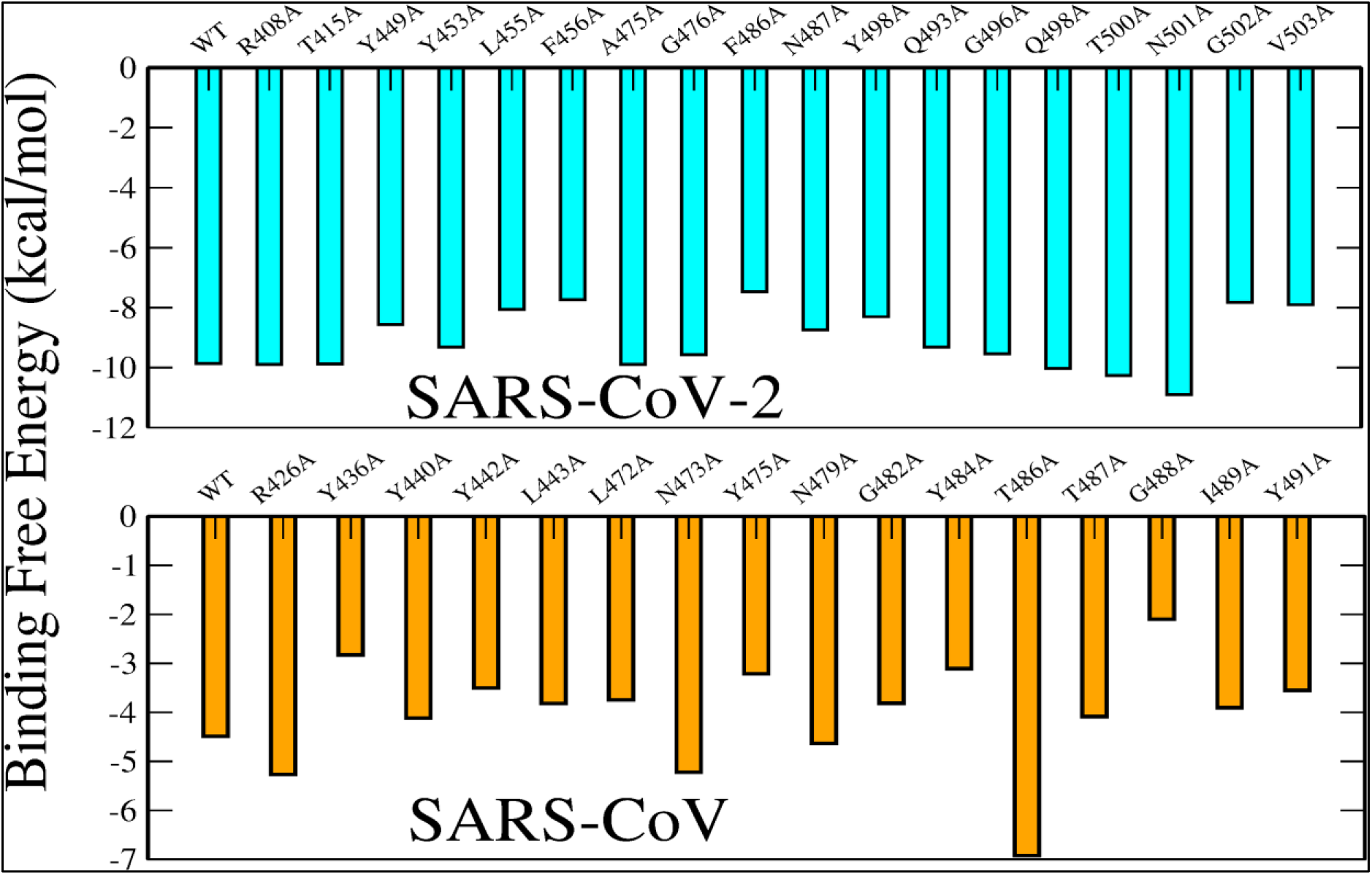
Computational alanine scanning results on SARS-CoV and SARS-CoV-2. The alanine scanning is performed on the residues of RBD domain that are coming in contact ACE2 receptor (cutoff value 4.0 Å).

Next section we discuss the results obtained from molecular dynamics simulations.

### 3.3 Energy components between RBD domains and hACE2 receptor: Hydrophobic interaction plays a major role

During 100 ns molecular dynamics simulations, both the adducts (SPIKE/hACE2) of SARS-CoV and SARS-CoV-2 are found to be stable. In **Figure S2** of Supporting Information, we have shown the root mean square deviation (RMSD) of Cα atoms ACE2 and RBD domains of SARS-CoV and SARS-CoV-2. From the RMSD plots and also visual inspections confirm the SPIKE/hACE2 complexes are fairly stable during the 100 ns simulations. Next section we compare the interaction energy and binding energy of SARS-CoV and SARS-CoV2.

From the molecular dynamics simulation trajectories, we extracted different components of interaction energy *i.e.* electrostatics and van-der-Waals (vdW) interaction terms between the RBD of SPIKE protein and the hACE2 receptor. In **Figure 4,** the probability of different components of interaction energies are shown between the SPIKE protein and the hACE2 receptor of SARS-CoV and SARS-CoV-2. The interaction energy plot reveals key features of how SARS-CoV and SARS-CoV-2 complexes gain stability. Overall, the RBD domain of SARS-CoV-2 shows slightly enhance interaction energy with hACE2 than SARS-CoV. The result is in good agreement with the recently published experimental results where it shows SPIKE protein on SARS-CoV-2 has a higher binding affinity toward hACE2 than SARS-CoV^6^. The increased interaction energy between SARS-CoV-2 is due to an increase in van-der-Waals interactions or the hydrophobic interactions between the RBD domain and the ACE2 receptor and there is a small decrease in electrostatic interactions. This also signifies that the substitution in the RBD domain facilitates the hydrophobic interactions, not the electrostatic interactions. One key hydrophobic residue Phe486 (Leu472 in SARS-CoV) which placed in a hydrophobic pocket created by the residues Leu79, Met82, and Tyr83 of hACE2 receptor. In the case of the SARS-CoV, the Leu472 is not exactly sitting in between the hydrophobic residues but juts oriented in outward-facing orientation (**Figure S2** of **Supporting Information**). This also signifies that the mutation that happened in SARS-CoV-2 increases the protein-protein hydrophobic contacts. This observation is very important as hydrophobic interaction may lead to a decrease in the number of water molecules near the receptor-binding domain. Also, it is a very common phenomenon found in many protein-protein and protein-DNA interfaces that water molecules play a crucial for stabilizing the binary complexes. A more in-depth analysis is given in **Section 3.4** where we discussed the role of interfacial water molecules in RBD/hACE2 complex.

**Figure 4.**
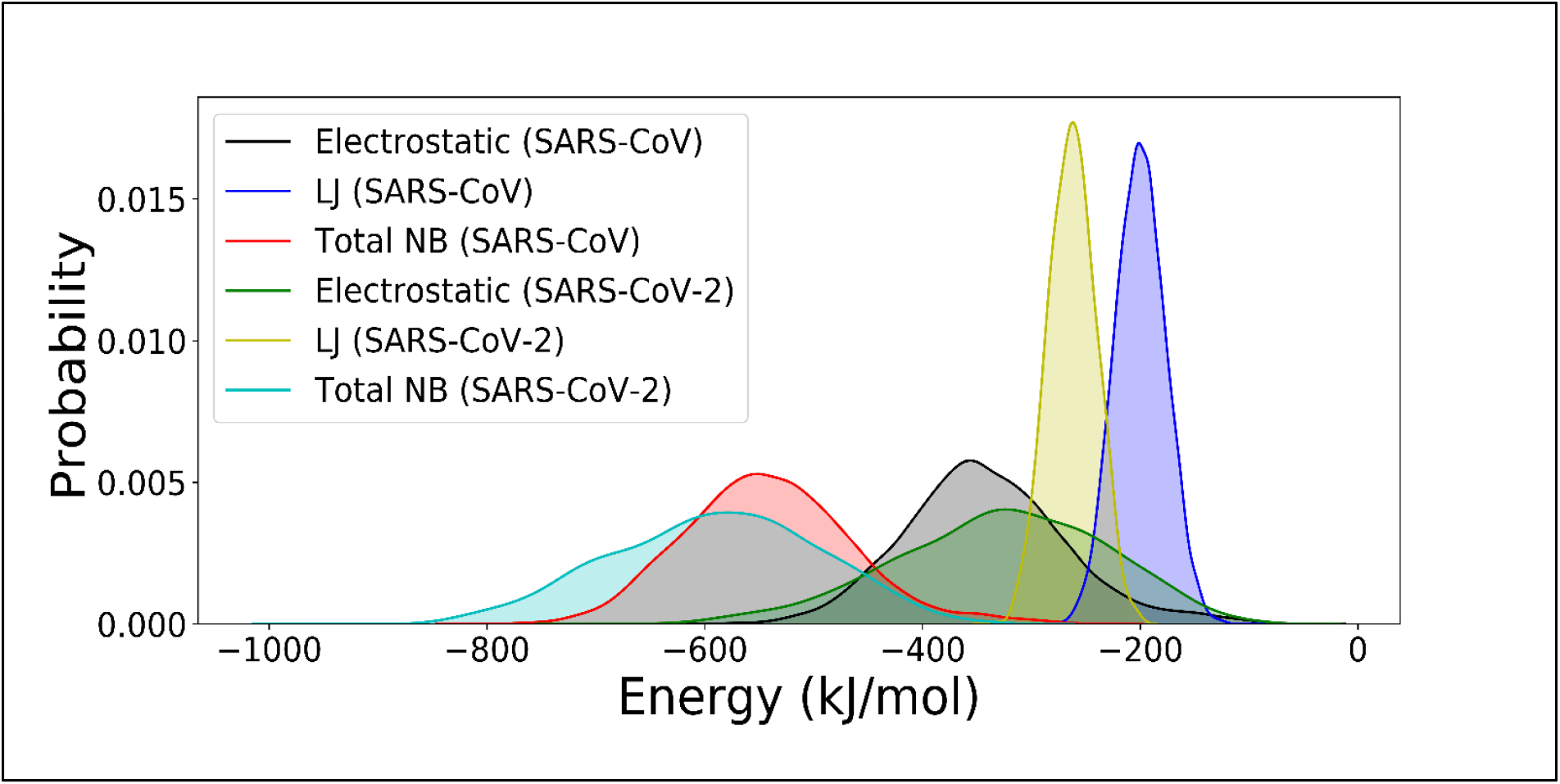
Different components of interaction energy between the RBD domains of SARS-CoV and SARS-CoV-2. Interaction energy is the sum of electrostatics and the van-der-Waals potential terms.

Earlier we computed the binding free energy for the crystal structures where SARS-CoV-2 has higher binding energy as compared to the SARS-CoV. Now, we calculate the binding affinity from simulation trajectories to see how the dynamics of these two domains affect the binding affinity. From 100 ns trajectory files, we extracted 1000 frames and calculate the binding free energy. The values that we got are −4.20± 0.62 kcal/ and −3.45±0.51 kcal/mol for SARS-CoV-2 and SARS-CoV. The dynamical picture obtained from the simulations signifies that the binding affinity of these two viruses is almost the same with hACE2 or slightly higher than SARS-CoV. Compared to the static picture, the dynamic picture indicates, fluctuation of both RBD and the hACE2 has significance on the overall binding affinity of the protein-protein interaction.

A key observation that was made from the simulation trajectories is that the minimum distance between the Cα atoms of RBD and the ACE2 domain varies significantly for SARS-CoV and SARS-CoV2 **(Figure 5 (a))**. The minimum Cα distance gives an idea of the relative separation between the two domains. It is observed that the average Cα distance for SARS-CoV is 5.16±02 Å and for SARS-CoV-2 the average distance between the two domains is 4.15±0.2 Å. The results signify that the SPIKE domain of SARS-CoV is staying ~1.0 Å far away from the hACE2 domain as that of SARS-CoV-2.

**Figure 5.**
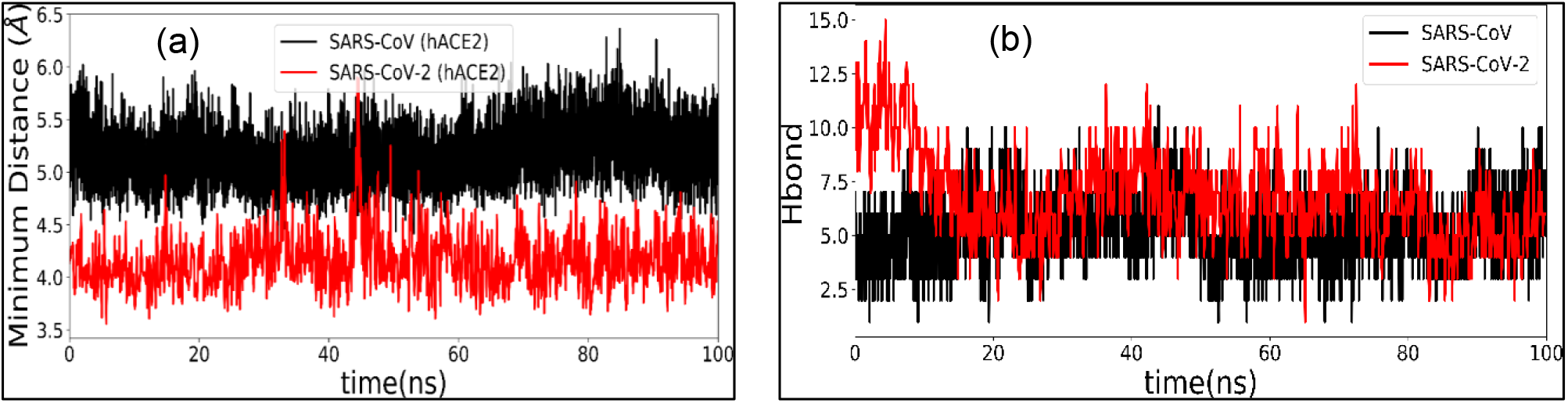
**(a)** Minimum distance between the Cα atoms of RBD and the hACE2 receptor during the MD simulations. **(b)** Number of hydrogen bonds between the RBD and the hACE2 receptor during the 100 ns MD simulations.

The reason for this may the poor binding of the RBD domain of SPIKE protein with hACE2. Besides, due to the substitution of four out of five proline-rich domain (472-483) for SARS-CoV-2 give more flexibility to the RBD that enhances the protein-protein interaction and the two domain remains in close proximity. Also, we have calculated the number of hydrogen bonds formed between the SPIKE domain and the hACE2 domain. In the crystal structure, there are 13 hydrogen bonds present in between SARS-CoV-2 (PDB ID: 6VWI), whereas it can be seen that the number of hydrogen bonds gradually decreases during the MD simulation and the average number of hydrogen bond is 6.7±2.1. In the case of SARS-CoV, the average hydrogen bond is 6 in the crystal structure whereas the average number of hydrogen bonds in the MD simulation is found to be 5.26±1.36 (see **Figure 5(b)**). This result was surprising, as the number of hydrogen bond is greatly reduced during the MD simulation for SARS-COV-2. Though the two domain remains in close proximity, the reduction in inter-domain hydrogen bonds signifies the enhanced hydrophobic interaction in SARS-CoV-2.

### 3.4 Role of Interfacial Water Molecules

It is well known that water-mediated interactions are one of the major driving forces of protein folding and protein-drug recognition processes^34–37^. In the previous section we observed the there is almost a 4-5 Å gap in between the SPIK/hACE2 protein-protein complex. There is a possibility that this interfacial domain or the gap region is hydrated with water molecules and those interfacial water molecules might play a significant role in stabilizing SPIKE protein and the hACE2 receptor.

Indeed, from our simulations, we observed that water molecules gradually populate the interfacial domain of SPIKE protein and hACE2 receptors. During the 100 ns simulation period, we observed that the interfacial domain forms a rich hydrogen-bonded network and stabilize the overall protein-protein complex.

From our simulations, we analyze the water molecules that come within the 3.5 Å near to the SPIKE protein and hACE2 domain. **Figure 6 (a)**, and (b) represent the snapshot of water molecules that are 3.5 Å away from the hACE2 and the RBD domain respectively. The snapshots are shown here only for the SARS-CoV-2, though a similar kind of results is also obtained for SARS-CoV.The fluctuation in the number of water molecules in the interfacial region of the ACE2 domain shows high water content for SARS-CoV than SARS-CoV-2 **(Figure 7(a)**). The average water molecules near the ACE2 domain are 214.2±8.68 and 242.7±10.04 for SARS-CoV-2 and SARS-CoV respectively Though the ACE2 interface is almost the same for SARS-CoV and SARS-CoV-2, there is a higher number of water molecules near the ACE2 interfaces. The probability plot (**Figure 7 (b)**) shows a clear difference in the number of water molecules between SARS-CoV and SARS-CoV-2. A similar result can be observed for the RBD domain of SPIKE protein. A greater number of water molecules can be seen near the SARS-CoV SPIKE domain than that of the SARS-CoV-2 SPIKE domain **(Figure 8(a) and (b)**). The high occupancy near the interfacial domain of SARS-CoV can be attributed to the fluctuation of the RBD and the hACE2 domain and larger gap in between these two domains.

**Figure 6.**
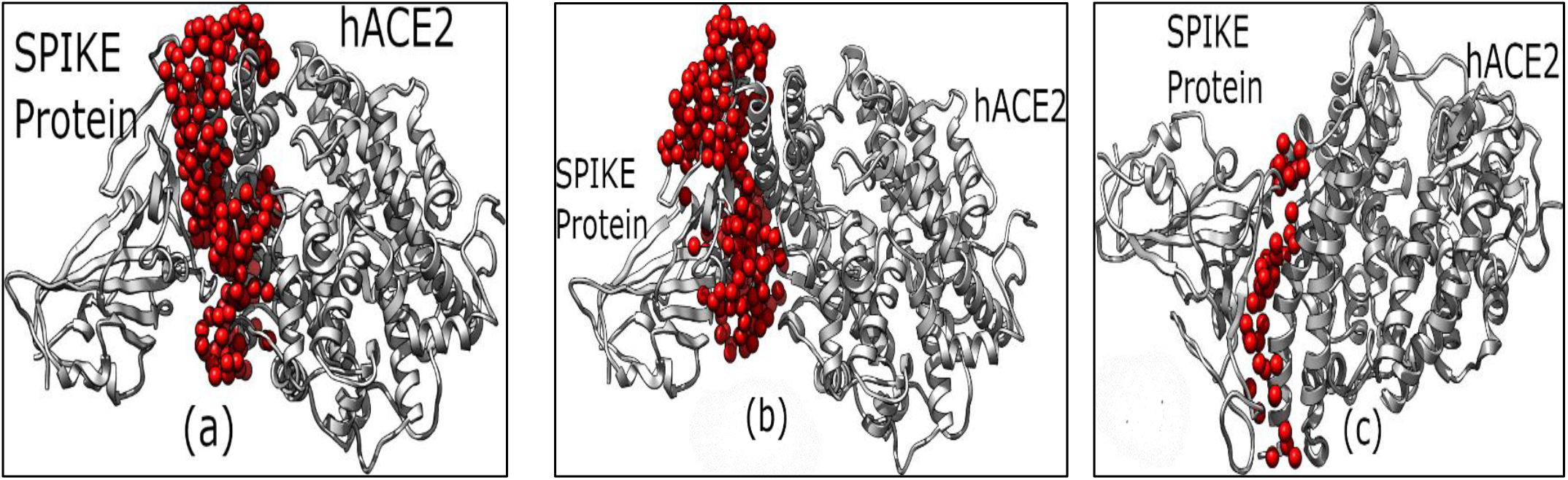
Snapshots of interfacial water molecules from molecular dynamics simulations. **(a)** Water molecules near 3.5 Å from the hACE2 domain. **(b)** Water molecules near 3.5 Å from the SPIKE-RBD domain, and **(c)** Bridge water molecules that are found 3.5 Å from both the RBD of SPIKE protein and hACE2 receptor. The above results are shown for the SARS-CoV-2. A similar result is obtained also for the SARS-CoV.

**Figure 7.**
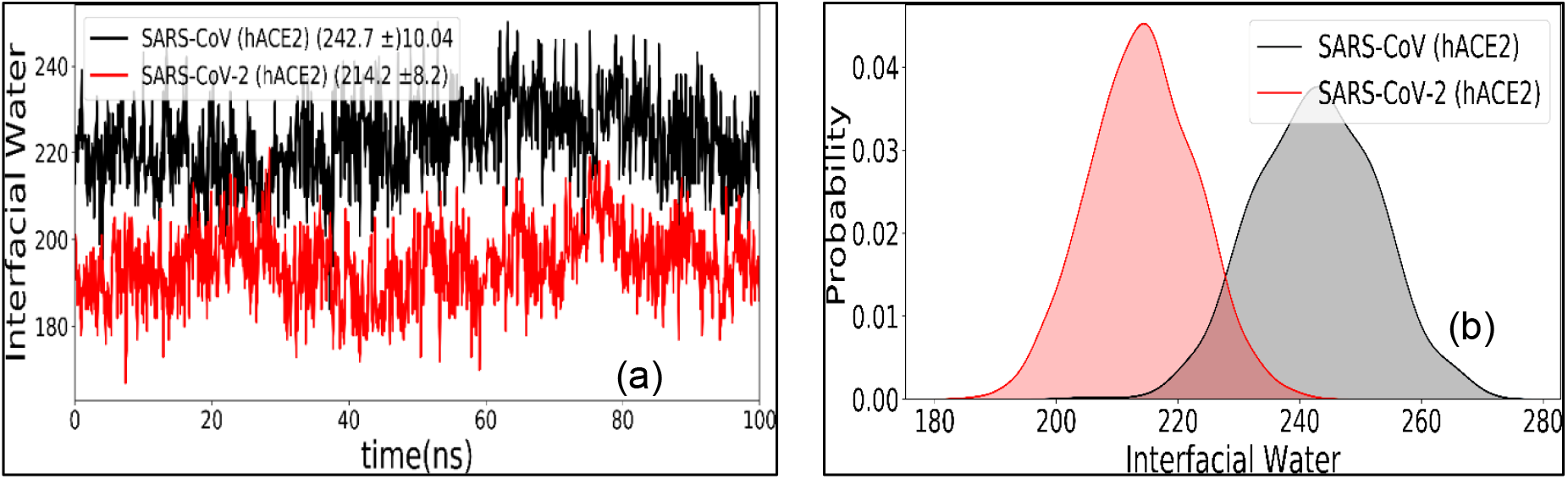
Interfacial water molecules near the hACE2 calculated from the 100 ns MD simulation trajectory files. Probabilty of interfacial water molecules that are near to the hACE2 surface.

**Figure 8.**
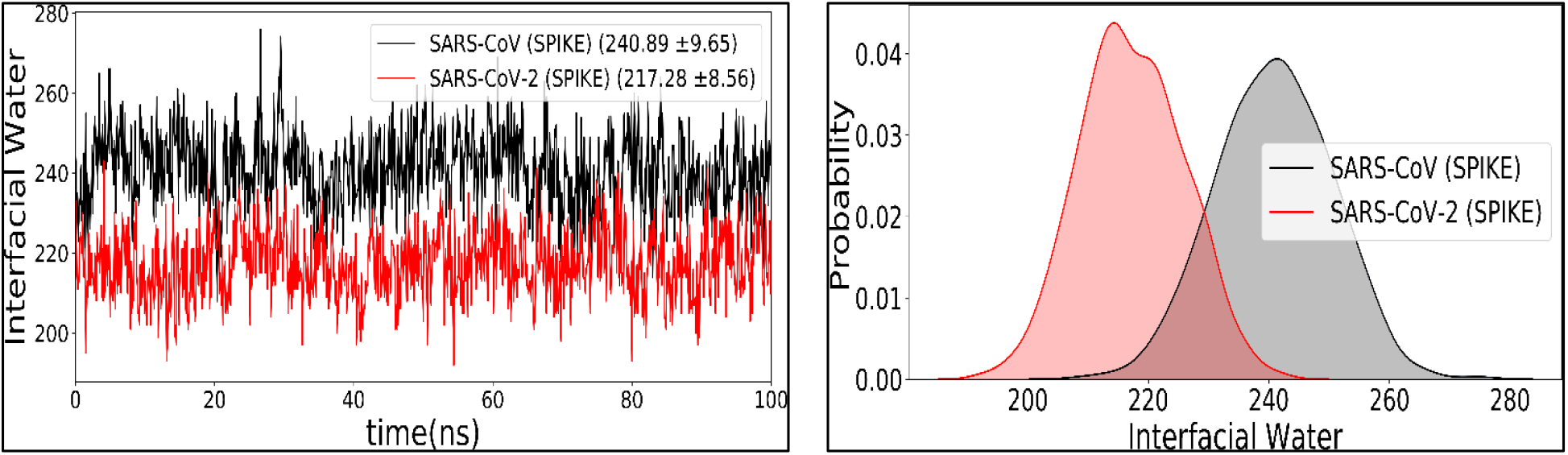
Interfacial water molecules near the RBD domain of SPIKE protein calculated from the 100 ns MD simulation trajectory files. Probabilty of interfcial water molecules that are near to the RBD surface.

Finally, we calculated the bridge-water molecules that are hydrogen-bonded with both ACE2 and RBD domains at the same time. We calculate the number of water molecules that are simultaneously forming hydrogen bonds with RBD and the ACE2 domain. In **Figure 9 (a)** we display the number of bridge water molecules during 100 ns molecular dynamics simulations and in **Figure 9 (b)** we show one bridge water molecule that is hydrogen-bonded to RBD (Gln493) as well as ACE2(Glu35) domain at the same time. From the simulation trajectories, several multi-water bridge water molecules can be observed in RBD and hACE2 domains. The bridge water molecules It can be seen that there is a slightly higher number of bridge water molecules (~2) for SARS-CoV-2 than that of SARS-CoV. These bridging water molecules play a significant role in stabilizing the two domains of the complex. We expect that these bridge water molecules play a significant role in stabilizing SPIKE/ACE2 domain. Here we want to highlight one key point that in the crystal structure (SARS-CoV-2) the Gln493 is engaged in a hydrogen bond with the Glu35, whereas from MD simulation we observed that water molecules enter into the interfacial region and connected these two residues.

**Figure 9.**
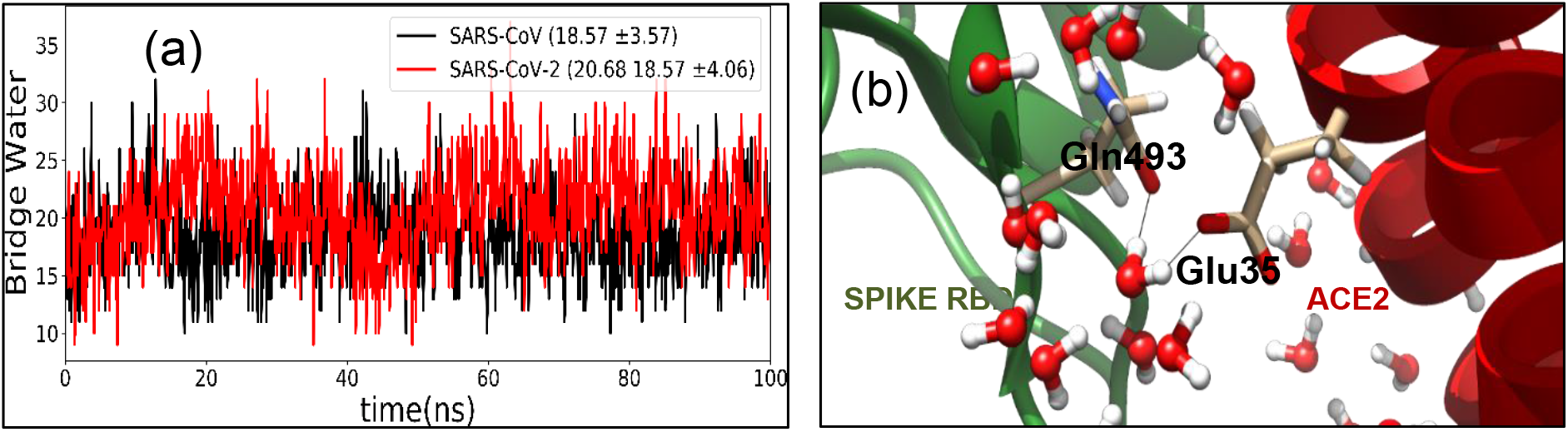
(a) Bridge Water molecules calculated from simulation trajectory files for SARS-CoV and SARS-CoV-2. (b) Representation of one of the bridge water molecules that hydrogen bonded with the both RBD and the hACE2 domain.

### 3.5 Energy Contribution from Interfacial Water Molecules

To have an idea about how the RBD and ACE2 domains are interacting with water molecules, we calculate the water-mediated hydrogen bonds with RBD and hACE2 domains. In the case of hACE2, the interfacial residues are making almost the same number of hydrogen bonds with the water molecules for SARS-CoV and SARS-CoV-2. This is expected as the ACE2 receptor is the same in both the viruses and small changes can be attributed to the fluctuation of the hACE2 domain. In **Figure 10 (a)**, the probability of the number of hydrogen bonds with hACE2 is shown. The average number of hydrogen bonds formed between the water molecules and the hACE2 domain are 113.9±6.4 and 113.2±6.9 for SARS-CoV and SARS-CoV-2 respectively. The significant changes can be seen near to RBD domain in between SARS-CoV and SARS-CoV-2 **(Figure 10(b)**). The average number of hydrogen bonds that are formed between the interfacial residues of RBD and water molecules are 115.02±6.4 and109.5±6.22 for SARS-CoV and SARS-CoV-2. The lesser number of hydrogen bonds for SARS-CoV-2 is the manifestation of mutation that occurred near the binding domain that enhances the hydrophobic nature at the interfacial domain.

**Figure 10.**
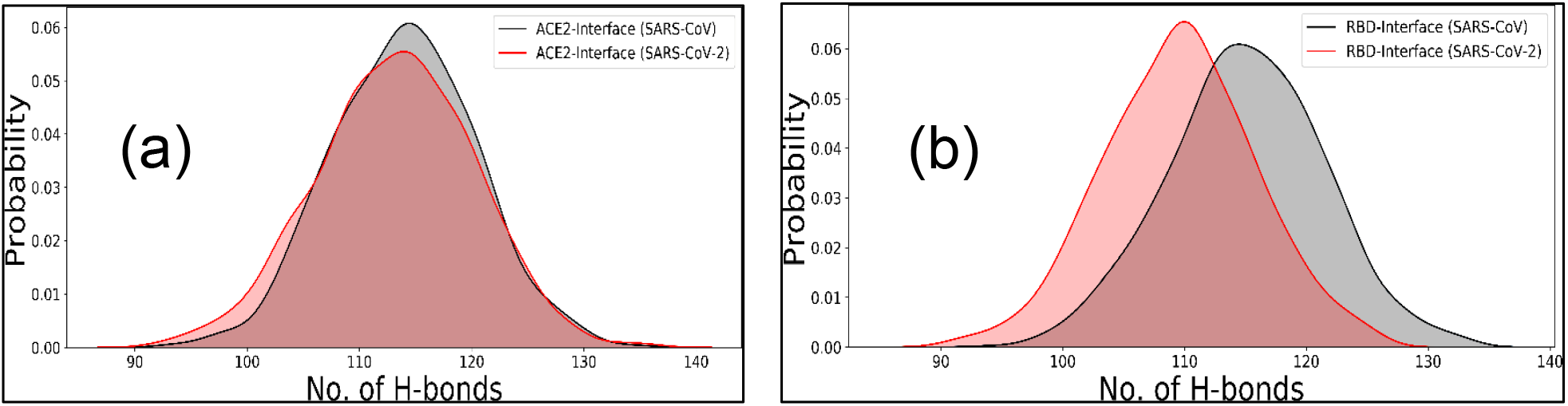
Probability of hydrogen binds between the water molecules and the interfacial domain of hACE2 and the RBD domains **(a)** For hACE2 receptor and **(b)** For RBD domain of SPIKE protein.

Finally, we calculated the interaction energy (electrostatic+ van der Waals) between the interfacial RBD and ACE2 domain with water molecules **(Figure 11(a),(b)**). We observe that the interaction energy between the water molecules and the hACE2 domain of both the viruses has almost the same that means the surface topology of hACE2 are behaves similarly during the MD simulations for both these viruses. The difference can be seen between the RBD of SARS-CoV and SARS-CoV-2. Interaction with water molecules is more preferred with SARS-CoV than SARS-CoV-2 as the water molecules make a greater number of hydrogen bonds with the RBD domain. Enhanced water interaction with the RBD of SARS-CoV reduces the direct interaction with the hACE2 receptor.

**Figure 11.**
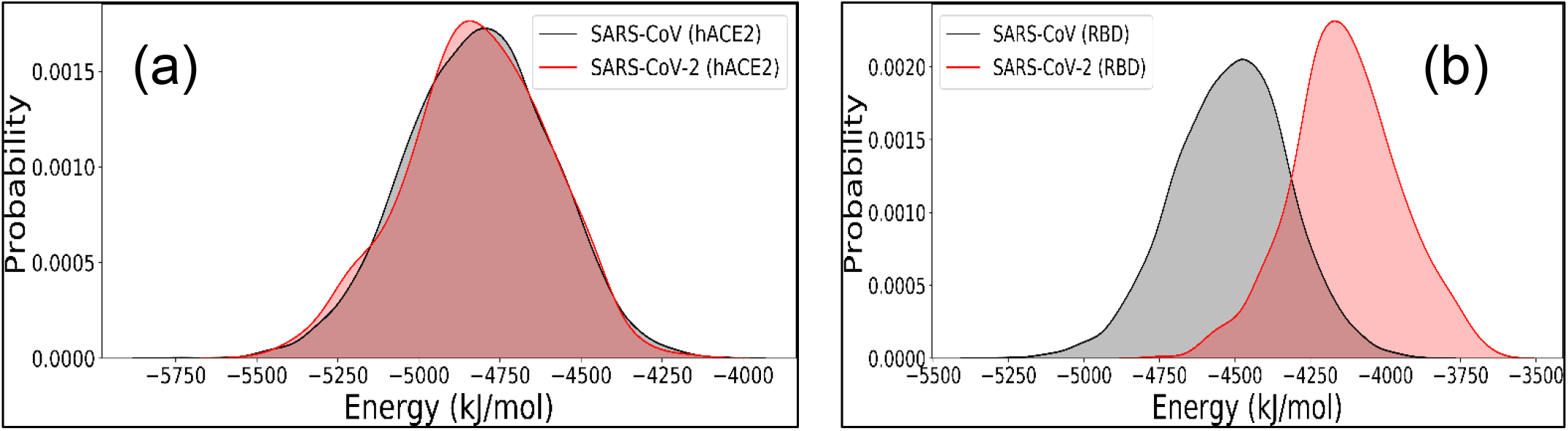
Total interaction energy (coulombic + van der Waals) between the water molecules and the hACE2 and the RBD domains.

## 4. Conclusion

In summary, we elucidate the similarities and differences between the SARS-CoV and SARS-CoV-2 by using molecular dynamics simulation approaches. From the structural point of view, the RBD of SPIKE protein shares a significant similarity in terms of the three-dimensional structure and the fold. However, significant differences are observed in how these two viruses bind to the hACE2 receptor. We found that improvement in hydrophobic contacts that leads to enhanced van der Waals interactions between the RBD and the hACE2 receptor affects the high binding affinity for the SARS-CoV-2. The most important information we found from the study is that involvement of water molecules at the interfacial domain of the RBD and ACE2 receptor. We observed that for both SARS-CoV and SARS-CoV-2 the interfacial domain is spontaneously hydrated and a significant number of water molecules populate the gap in between RBD and ACE2 domain. It is found that bridge water molecules play a significant role in stabilizing the protein-protein binary complex. This kind of water-mediated interaction for SARS-CoV and SARS-CoV-2 did not observe in the previous study. The present study unravels the many structural and dynamical features of this protein-protein dimer complexes that help us to understand how this virus interacts with the human cell.

