## Supporting Information for "Interfacial Water Molecules Make RBD of SPIKE Protein and Human ACE2 to Stick Together"

Quantumzyme LLP, Bangalore India 560027

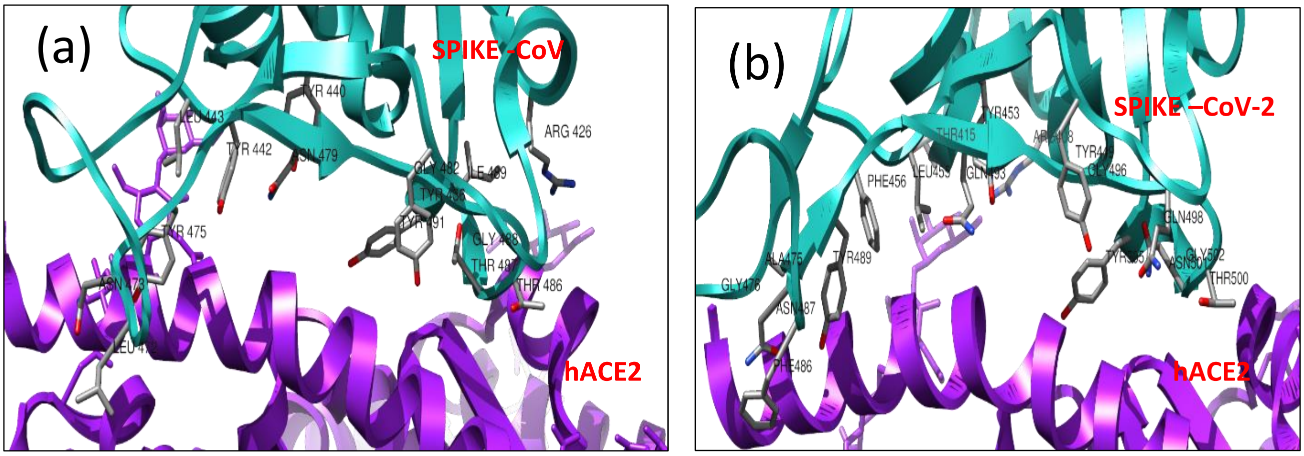


**Figure S1 (a) SARS-CoV:** The interfacial residues of RBD domain of SPIKE protein near to the 4 Å of hACE2 receptor. (b) **SARS-CoV-2:** The interfacial residues of RBD domain of SPIKE protein near to the 4 Å of hACE2 receptor.

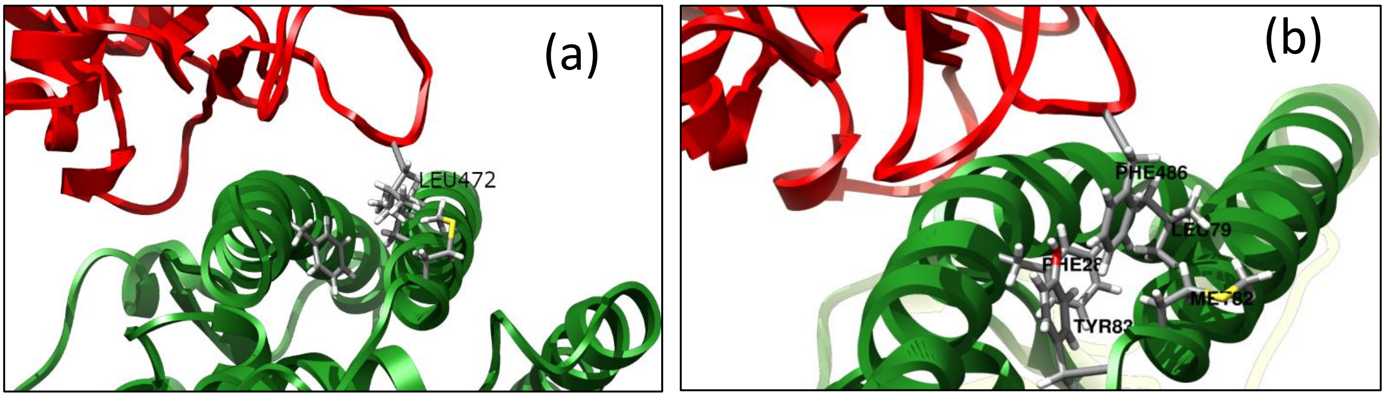


**Figure S2** Average position found for Leu472 (SARS-CoV) and Phe486 (SARS-CoV-2) from the molecular dynamics simulation trajectories.

| Residues | SARS-CoV-RBD (4Å) | ACE2 (4Å) | SARS-COV-2-RBD (4Å) | ACE2 (4Å) | SARS-CoV-RBD (8Å) | ACE2 (8Å) | SARS-COV-2-RBD (8Å) | ACE2 (8Å) |
|---|---|---|---|---|---|---|---|---|
| 1 | Y491 | R357 | Y505 | R393 | Y494 | V506 | Q506 | R393 |
| 2 | I489 | D355 | G502 | R357 | Q492 | H505 | Y505 | F390 |
| 3 | G488 | G354 | N501 | D355 | Y491 | F504 | G504 | Q388 |
| 4 | T487 | K353 | T500 | G354 | G490 | L503 | V503 | A387 |
| 5 | T486 | N330 | Q498 | K353 | I489 | R393 | G502 | A386 |
| 6 | Y484 | E329 | G496 | N330 | G488 | Q388 | N501 | R357 |
| 7 | G482 | Q325 | Q493 | Y83 | T487 | A387 | T500 | F356 |
| 8 | N479 | Y83 | Y489 | M82 | T486 | A386 | P499 | D355 |
| 9 | Y475 | M82 | N487 | L79 | Y485 | R357 | Q498 | G354 |
| 10 | N473 | L79 | F486 | L45 | Y484 | F356 | F497 | K353 |
| 11 | L472 | L45 | G476 | Q42 | F483 | D355 | G496 | G352 |
| 12 | L443 | Q42 | A475 | Y41 | G482 | G354 | Y495 | L351 |
| 13 | Y442 | Y41 | F456 | D38 | Y481 | K353 | S494 | N330 |
| 14 | Y440 | D38 | L455 | E37 | D480 | G352 | Q493 | E329 |
| 15 | Y436 | E37 | Y453 | E35 | N479 | L351 | L492 | F327 |
| 16 | R426 | H34 | Y449 | H34 | W476 | N330 | P491 | G326 |
| 17 | | K31 | T415 | K31 | Y475 | E329 | F490 | Q325 |
| 18 | | F28 | R408 | F28 | C474 | F327 | Y489 | T324 |
| 19 | | T27 | | T27 | N473 | G326 | C488 | P84 |
| 20 | | Q24 | | Q24 | L472 | Q325 | Q487 | Y83 |
| 21 | | | | S19 | A471 | T324 | F486 | M82 |
| 22 | | | | | P470 | Y83 | G485 | Q81 |
| 23 | | | | | K465 | M82 | E484 | A80 |
| 24 | | | | | G464 | A80 | T478 | L79 |
| 25 | | | | | E463 | L79 | S477 | T78 |
| 26 | | | | | P462 | T78 | G476 | Q76 |
| 27 | | | | | S461 | Q76 | A475 | K68 |
| 28 | | | | | F460 | E75 | Q474 | W48 |
| 29 | | | | | K447 | N49 | Y473 | L45 |
| 30 | | | | | H445 | W48 | F456 | Q42 |
| 31 | | | | | R444 | L45 | L455 | Y41 |
| 32 | | | | | L443 | S43 | R454 | L39 |
| 33 | | | | | Y442 | Q42 | Y453 | D38 |
| 34 | | | | | R441 | Y41 | Y449 | E37 |
| 35 | | | | | Y440 | L39 | Q448 | A36 |
| 36 | | | | | Y436 | D38 | G447 | E35 |
| 37 | | | | | N435 | E37 | T446 | H34 |
| 38 | | | | | G434 | E35 | S445 | N33 |
| 39 | | | | | T433 | H34 | R439 | F32 |
| 40 | | | | | S432 | N33 | Y421 | K31 |
| 41 | | | | | T431 | F32 | D420 | D30 |
| 42 | | | | | R426 | K31 | V417 | L29 |
| 43 | | | | | N424 | D30 | G416 | F28 |
| 44 | | | | | Y408 | L29 | T415 | T27 |

| | | | | | | | | |
|---|---|---|---|---|---|---|---|---|
| 45 | | | | | D407 | F28 | Q414 | K26 |
| 46 | | | | | V404 | T27 | G413 | A25 |
| 47 | | | | | G403 | K26 | Q409 | Q24 |
| 48 | | | | | T402 | A25 | R408 | E23 |
| 49 | | | | | Q401 | Q24 | D406 | I21 |
| 50 | | | | | R395 | E23 | D405 | T20 |
| 51 | | | | | D393 | I21 | K403 | S19 |
| 52 | | | | | D392 | T20 | | |
| 53 | | | | | K390 | S19 | | |

**Table S1** Contact residues RBD domain and hACE2 receptor for SARS-CoV and SARS-CoV-2 using the cut-off value of 4 Å and 8 Å.

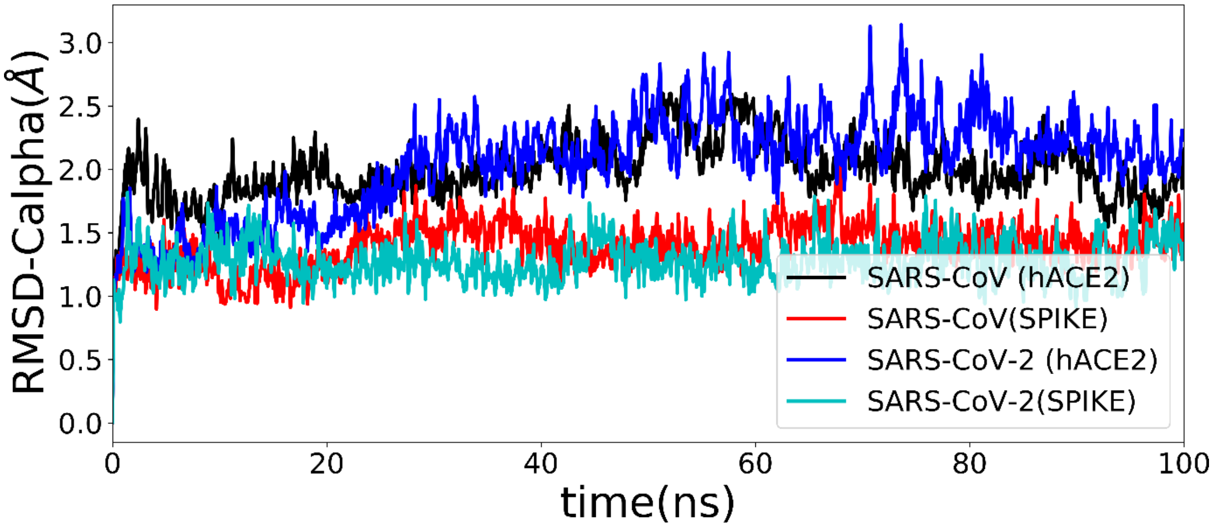


**Figure S2** Fluctuation of RMSD of Cα atoms of SPIKE protein and hACE2 receptor for SARS-CoV and SARS-CoV-2 during 100 ns molecular dynamics simulations.
